# Alternative Methods for Mitochondrial Transplantation: Efficiency of Unpackaged and Lipid-Packaged Preparations

**DOI:** 10.1101/102913

**Authors:** DL McPhie, LW Sargent, SM Babb, D Ben-Shachar, AM Cataldo, BM Cohen

## Abstract

Mitochondrial transplantation is currently being explored as a means to repair and restore proper organelle function in a variety of inherited and acquired disorders of energy metabolism. The optimal preparation and application of donor mitochondria is unknown, but most studies *in vivo* have used injection techniques or, for tissue studies, unpackaged mitochondria (organelles isolated and suspended in buffer) in transplant experiments. Packaging in lipid rafts can increase recipient cell uptake of some compounds and objects. We present the first data comparing recipient cell uptake of unpackaged mitochondria to recipient cell uptake of mitochondria packaged in cell membrane lipids. Mitochondria and membranes were prepared from autologous cells and applied to cells (fibroblasts) in culture. Both unpackaged and lipid-packaged mitochondria were taken into recipient cells and the donor mitochondria showed evidence, in each case, of retained functionality and the ability <u>to merge with the recipient mitochondrial matrix</u>. However, lipid packaging appeared to enhance the uptake of functional mitochondria. Current studies of mitochondrial transplantation in animal models might fruitfully explore the utility and efficacy of lipid-packaged mitochondria in transplant experiments.

## Introduction

Mitochondria are essential organelles in all eukaryotic cells. They appear to have originated from engulfed prokaryotes, and they play a key role in bioenergetic processes, especially oxidative phosphorylation and the aerobic metabolism of glucose and fat. They also play a central role in calcium storage and signaling and in apoptosis (1,2). The human mitochondrial genome is maternally inherited, small, at 16,568 bp, and encodes only a limited number of mitochondria-specific proteins, rRNAs, and tRNAs (3). All other mitochondrial proteins are encoded by the nuclear genome. Notably, the mitochondrial genome undergoes a high rate of mutation because mtDNA is not protected by histones, is inefficiently repaired (4) and is exposed to oxygen radicals generated by oxidative phosphorylation (1).

A large number of heritable diseases are caused by mutations in mitochondrial and nuclear genes encoding mitochondrial proteins. Many produce heritable neurologic syndromes and skeletal or cardiac myopathies (1,3,5,6). In addition, environmentally induced mutations in mtDNA have been implicated in commonly acquired disorders, including ischemic diseases of the heart and brain, neurodegenerative diseases, some liver diseases, and some cancers (5,7). Mitochondrial compromise appears especially relevant to normal brain and muscle function, as these tissues use far more energy per unit weight than other tissues of the body. Thus, mitochondrial dysfunctions are often observed in neuropsychiatric disease, including bipolar disorders (BD), depressive disorders, schizophrenias, and Rett’s syndrome; neurodegenerative disease like Alzheimer’s disease, Parkinson’s disease, Friedeich’s ataxia (and other ataxias); amyotrophic lateral sclerosis (ALS) (and other motor neuron diseases); Huntington’s disease; and various neuropathies and myopathies, such as Leber’s hereditary optic neuropathy (LHON), encephalopathies, lactic acidosis, stroke syndromes (MELAS); macular degeneration; epilepsies; and mitochondrial myopathies. Of great importance, mitochondrial structural integrity and functional efficiency declines with age, and mitochondrial alterations may be a key determinant of age related reductions in tissue health, even in the absence of pathology (8).

Mitochondrial networks are highly dynamic (9). Within cells, the networks expand and contract, in response to local signals. Through budding and fission, new mitochondria leave the local network. Simultaneously, fusion of existing mitochondria occurs between contiguous networks (10). These activities suggest that mitochondrial pathologies might be addressed by altering mitochondrial growth and connectivity or by replacing damaged mitochondria (11).

Pharmacologic and nutritional precursor approaches to enhance mitochondrial function and biogenesis are being studied, but have not yet produced successful treatment strategies (11,12).

In *in vitro* cell culture, intercellular transfer of functionally competent mitochondria has been observed, and this mechanism may improve local mitochondrial deficiencies or compensate for defective mitochondria (13). Mimicking this natural mechanism, the replacement or transplant of compromised mitochondria with healthy exogenous organelles might have therapeutic potential. We have proposed techniques and uses for such replacement therapies (14) and there have been increasing numbers of reports of mitochondrial transplantation in both *in vitro* and *in vivo* studies. Transplantation methods vary by study, but include direct injections of mitochondria and various techniques to induce fusion of cells to allow transfer of mitochondria through cell membranes (Reviewed by McCully, 2016 (15)). One technique, not yet reported, involves the use of lipid rafts. These have been used elsewhere to induce cells to engulf exogenous substances and organelles. We hypothesized that this packaging could be useful to enhance mitochondrial transplantation or uptake, as well.

In this study, we compare the efficiency of recipient cell uptake of mitochondria following the delivery of donor mitochondria in buffer solution to donor mitochondria mixed with lipid rafts derived from the recipient cells’ outer membrane. Our hypothesis was that preparations mixed with lipid membranes would achieve better incorporation into recipient cells. Higher incorporation of transplanted mitochondria might be important for achieving therapeutic benefits.

## Materials and methods

### Cell culture

Human fibroblasts were established from skin biopsy samples from study participants at McLean Hospital (Belmont, MA). Study procedures were approved by the McLean Hospital Institutional Review Board, and all subjects gave written consent to participate after procedures and possible side effects were explained. Fibroblasts were expanded in cell culture media (Minimal Essential Medium, NEAA (Gibco, ThermoFisher) supplemented with 15% fetal bovine serum, 1% GlutaMAX (Gibco, ThermoFisher), and 1% Penicillin-streptomycin in 5% CO_2_ at 37**°**C. Media was changed every other day. 1X TrypLE Express Enzyme (Gibco, ThermoFisher) was used for passaging cells. Cell lines used for experiments were all between passages P5 and P7.

### Isolation of mitochondria

Mitochondria were isolated from fibroblasts using a modified protocol from Frezza et al., 2007 (16). Ten mL of IB_C_ buffer (1 mL of 0.1 M Tris/MOPS, 100 uL of 0.1 M EGTA/Tris, and 2 mL of 1 M sucrose diluted with distilled water, pH 7.4) was prepared and kept on ice. Two days prior to the experiment, cells were plated to reach 80-85% confluence on the day of the experiment. On the day of the experiment, cells were washed twice with DPBS (without Ca^2+^ and Mg^2+^) and detached using 1X TrypLE Express Enzyme. Cells were resuspended in cell culture media and transferred to a new conical tube, centrifuged for three minutes at 600 g and then counted. 2 x 10^7^ fibroblasts were transferred to a separate tube and centrifuged for ten minutes at 600 g. The supernatant was discarded and the cell pellet was resuspended in IB_C_ buffer. Cells were then homogenized using a 22G needle until at least 50% of the cells were damaged (determined by visual check under a microscope), then the resulting homogenate was centrifuged for ten minutes at 600 g at 4**°**C. The supernatant was collected, transferred to a pre-chilled microcentrifuge tube, and centrifuged for ten minutes at 7,000 g at 4**°**C. The supernatant was discarded and the pellet was resuspended in IB_C_ buffer. The pellet suspension was centrifuged again for ten minutes at 7,000 g at 4**°**C. The supernatant was discarded and the pellet containing the isolated mitochondria was carefully resuspended in IB_C_ buffer to avoid the formation of bubbles and kept on ice.

### Assessing the purity and membrane intactness of mitochondrial isolates

Donor mitochondria were isolated, pelleted, and fixed in 2% glutaraldehyde, 100 mM sodium cacodylate buffer, 2 mM CaCl_2_ (pH 7.4). Mitochondrial pellets were processed following standard electron microscopy protocols. All chemicals were from Ted Pella, Inc. (Redding, CA). Briefly, mitochondrial pellets were postfixed in 1% OsO_4_, dehydrated in a series of alcohols up to absolute alcohol and then transitioned to propylene oxide and embedded in Epon. Blocks were sectioned, stained, and examined on a JEOL 1200EX microscope. Electron microscopy was used to determine the presence of cristae, and the presence of the inner and outer membranes of the mitochondria as an indication of the intactness of mitochondrial membranes. Western blots of isolated mitochondria were also run and probed with an antibody to TOM-20, a specific marker protein for mitochondrial outer membrane, in order to test for enrichment and confirm intactness of mitochondria in the mitochondrial isolates.

### Assessing the functional status of mitochondrial preparations

The functional status of isolated mitochondria was assessed by incubating an aliquot of the mitochondrial preparation with the dye JC-1 following manufacturer’s (Life Technologies) protocols. The aliquot was visualized under a fluorescent microscope using a filter set with 488nm excitation and 510nm emission for the green channel and a filter set with a 549nm excitation and 575nm emission for the red channel. JC-1 is an indicator of mitochondrial membrane potential. In healthy mitochondria, JC-1 accumulates and forms aggregates that fluoresce red, while at depolarized membrane potentials, JC-1 exists as a green-fluorescent monomer.

### Recipient fibroblasts

Two days prior to the experiment, recipient fibroblasts were plated at 2.1 x 10^4^ cells per well in all four wells of Lab-Tek II chamber coverglasses (Nunc) coated with fibronectin (5 ug/mL final concentration).

### Labeling donor and recipient fibroblast mitochondria

Sixteen hours prior to the experiment, donor fibroblasts were infected with either HSV-DsRed2-mito or HSV-Cy-5-mito, which use the mitochondrial targeting sequence of the human cytochrome oxidase subunit VIII to target mitochondria and cause the expression of either a red fluorescent protein (DsRed2) or a far-red fluorescent protein (Cy-5) (HSV constructs and virus prepared by R. Neve, Viral Core Facility McGovern Institute for Brain Research, MIT). Infection was achieved by adding the appropriate virus at a multiplicity of infection (MOI) =1 directly to donor cells in culture. Recipient fibroblast mitochondrial networks were labeled with CellLight Mitochondria-GFP, BacMam 2.0 following the manufacturer’s (ThermoFisher) protocol. CellLight Mitochondria-GFP uses the leader sequence of E1 alpha-pyruvate dehydrogenase to specifically target mitochondria and cause the expression of green fluorescent protein (GFP). Both donor and recipient fibroblasts were incubated overnight in 5% CO_2_ at 37**°**C. On the day of the experiment, recipient fibroblasts were also labeled with Hoechst 33342 (1:100) to stain the nuclei.

### Preparation of Detergent-free lipid rafts

Detergent-free lipid rafts were isolated from fibroblasts using a modified protocol from Macdonald and Pike (17). All procedures were done on ice. Fibroblasts were plated two days prior to the experiment. On the day of study, cells from four nearly confluent 15 cm plates were scraped into base buffer (20 mM Tris HCl, pH 7.8, 250 mM sucrose) supplemented with 1 mM CaCl_2_ and 1 mM MgCl_2_. Cells were pelleted by centrifugation for two minutes at 250 g, then resuspended in base buffer supplemented with 1 mM CaCl_2_, 1 mM MgCl_2_, and protease inhibitors (100 mM PMSF, 1 M β glycerol PO_4_, 1 M NA_3_VO_4_, 10 ug/ml aprotinin, 4 mg/mL leupeptin, 1 mM benzamidine, 0.5 M NEM, 1 M NAF). Cells were lysed using a 22G needle, then centrifuged for ten minutes at 1000g. The resulting supernatant was collected and transferred to a new tube. Additional base buffer supplemented with 1 mM CaCl_2_, 1 mM MgCl_2_, and protease inhibitors was added to the remaining pellet. Shearing with a 22G needle and subsequent centrifugation were repeated once and the resulting two supernatants were combined. An equal volume of 50% OptiPrep Density Gradient (Sigma) was added to the combined supernatants and this 25% lysate was transferred to a centrifuge tube. A step gradient of 0% to 20% OptiPrep, diluted in base buffer including 1 mM CaCl_2_ and 1 mM MgCl_2,_ was layered on top of this mixture. The gradient was centrifuged for 90 minutes at 52,000g using an SW-41 rotor in a Beckman ultracentrifuge. After centrifugation, a distinct band containing the lipid rafts, visible between the 20% and 25% layers, was removed using a 20G needle, transferred to a fresh microcentrifuge tube, and kept on ice.

### Mitochondrial transplantation

Donor mitochondria (expressing DsRed2) with and without lipid rafts were transplanted into autologous recipient fibroblasts. To verify that there was no cross contamination, two side-by side control conditions were also tested: rafts only and recipient cells only. Immediately prior to transplantation, mitochondria isolated from donor cells were gently resuspended in 100ul of imaging media (MEM without phenol red and glutamine, plus 10% fetal bovine serum, and 1% GlutaMax) and divided into two samples to be prepared, with and without lipid rafts. The four conditions to be tested were prepared as follows: a) Mito plus rafts: 50ul mitochondrial isolate was mixed with 200ul of lipid rafts and then an additional 100ul of imaging media was added, b) Mito without rafts: 50ul of mitochondrial isolate was mixed with 200ul of imaging media and then another 100ul of imaging media was added, c) Rafts only: 200ul of lipid rafts was mixed with 150ul of imaging media, d) Recipient cells only: 350 ul of imaging media. Media was removed from the wells containing recipient fibroblasts, which had been previously labeled with CellLight Mitochondria GFP. Then, donor mitochondrial preparations were added drop-wise to the wells containing the recipient fibroblasts. Cells were incubated for twenty minutes at 37**°**C then washed once with imaging media to remove residual mitochondria not taken up by recipient cells. All conditions were then fixed with paraformaldehyde (4% paraformaldehyde in 100 mM phosphate buffer, pH 7.4) for fluorescent imaging.

### Imaging mitochondria in recipient fibroblasts

After incubation of the preparations, as described above, cells were fixed in 4% paraformaldehyde in 100 mM phosphate buffer, pH 7.4, and mitochondrial uptake was assessed using a fluorescent Zeiss Observer Z.1 microscope. Donor mitochondria labeled with DsRed2 were visualized at excitation (549 nm) and emission (575 nm). Recipient mitochondria labeled with Mitochondria-GFP were visualized at excitation(488 nm) and emission (510 nm). Recipient nuclei were visualized at excitation and emission of 350 nm and 461 nm. Images were taken at 40X. Additional studies using confocal microscopy were done to assess the results of a randomly chosen sub-sample of the transplantations. Confocal microscopy was performed on a laser scanning confocal microscope (SP8-TCS, Leica). Eight Z stack images, with Z section intervals of 0.75µm (using a 40X 1.3NA lens) of fields of cells were acquired at excitation wavelengths of 405, 488 and 561 nm. This Z section interval, somewhat smaller than the average mitochondrial diameter, provided adequate resolution to observe whether your recipient and donor mitochondria remain physically separate.

### Quantification of donor recipient cells taking up donor mitochondria

Ten fields taken from the middle of confocal Z stacks showing approximately twenty cells each were examined. Cells were classified as having taken up mitochondria if both recipient (labeled with GFP) and donor mitochondria (labeled with DsRed2) were visible within the same cell.

### Live imaging of internalized mitochondria for assessment of mitochondrial function

Donor mitochondria were infected with HSV-Cy5-mito, a far-red fluorescent dye, then isolated and incubated for 20 minutes with recipient cells that had been labeled with mitochondrial-GFP (green) as outlined above. Prior to imaging, fibroblasts treated with mitochondrial preparations were incubated with tetramethylrhodamine, ethyl ester (TMRE) (100nM), an orange fluorescent dye that is readily taken up by active mitochondria. Live imaging was done using fluorescent Zeiss Observer Z.1 at 40X. Healthy mitochondria, labeled with TMRE, were visualized at excitation and emission of 549 nm and 575 nm, respectively. Donor mitochondria were visualized at excitation and emission of 647nm and 665 nm. Recipient mitochondria were visualized at excitation and emission of 488 nm and 510 nm, respectively. Recipient cell nuclei were visualized at excitation and emission of 350 nm and 461 nm, respectively.

## Results and Discussion

Unpackaged mitochondria and lipid-packaged mitochondria were prepared in parallel and recipient cells were exposed at the same time, to enhance accuracy of comparisons. Figure 1 shows that the isolated mitochondria for donor preparations were structurally intact and functionally competent, as evidenced by red JC-1 staining and the presence of intact cristae and two membranes on the EM level (B). Additionally, western blots of the mitochondrial isolates show an increased signal for TOM-20 indicating mitochondrial enrichment in the isolates versus unprepared lysates (data not shown).

**Figure 1.**
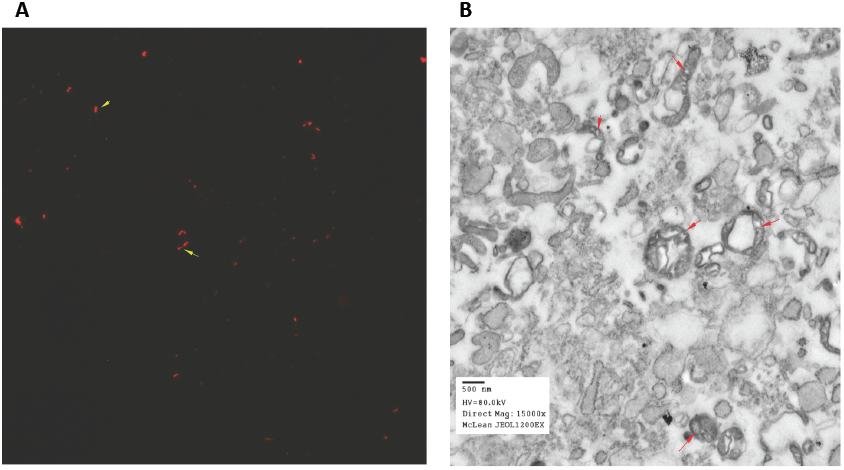
Isolated mitochondria appear healthy on light and electron microscopy. A) Isolated mitochondria stained with JC-1; JC-1 aggregates in healthy mitochondria,as evidenced by a red (575nm) fluorescence. (B) Electron micrograph of isolated mitochondrial pellet showing intact mitochondria as evidenced by intact cristae.

Uptake of mitochondria by recipient cells was observed in both unpackaged mitochondria and mitochondria packaged in lipid rafts (Figure 2). Both red donor mitochondria and green recipient mitochondria are visible as well as some instances of potential mitochondrial fusion (yellow). Most mitochondria exist in networks, and the observed fusion may be between mitochondrial networks, rather than isolated mitochondria.

**Figure 2.**
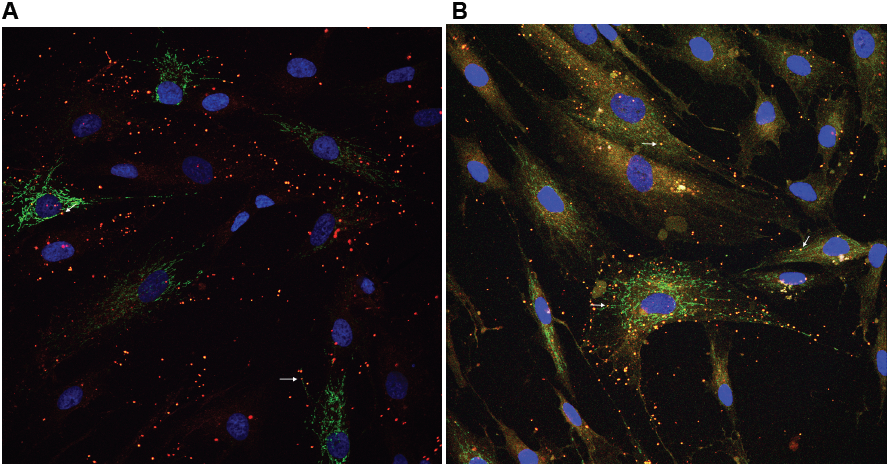
Recipient cells take up donor mitochondria that are unpackaged (A) or packaged in lipid rafts (B). Each case shows a single slice from the middle of a Z stack taken by confocal microscopy showing isolated mitochondria (red) at the same levels as recipient mitochondrial networks (green). Fibroblast nuclei appear blue. Arrows indicate a few examples of points of potential fusion between donor and recipient mitochondria. Potential fusion appears as yellow areas, where donor (red) mitochondria co-occur with recipient (green) mitochondria. In both cases, naked or lipid-packaged, mitochondria appear to enter recipient cells.

Relative efficiency of uptake is shown in Figure 3. Given similar amounts of donor mitochondria, recipient cells took up significantly more if the mitochondria were packaged in lipids. Almost twice as many cells took up mitochondria packaged in lipids (mito+rafts) compared to unpackaged mitochondria (mito only). Even at 1/10th the concentration of unpackaged mitochondria, over half as many recipient cells took up the donor mitochondria packaged in lipids (1/10 mito+rafts), as those suspended in buffer alone. These differences were statistically significant.

**Figure 3.**
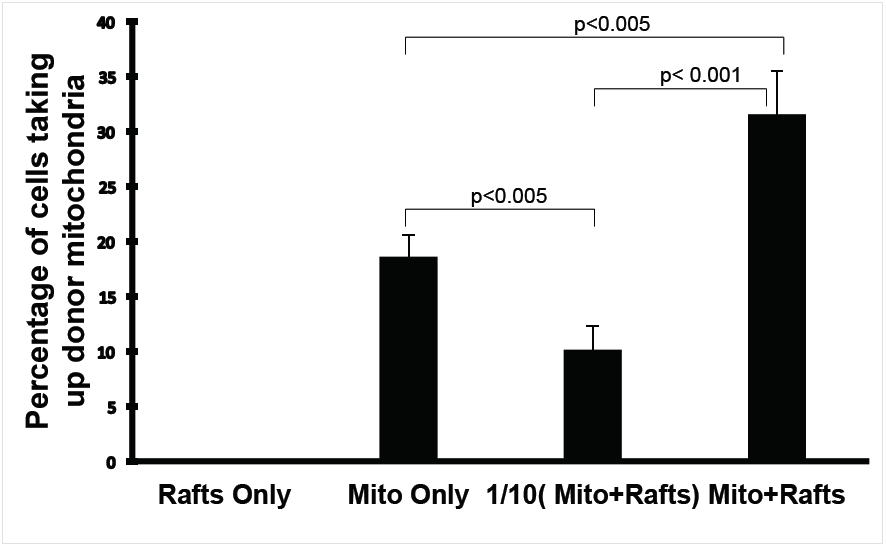
Quantification of the efficacy of the uptake of naked mitochondria versus mitochondria mixed with lipid rafts. Efficiency of uptake of mitochondria was determined by counting the number of cells that had taken up mitochondria, as observed by confocal microscopy. Ten random fields per condition were used for counting. Data are shown as percentage of total cells per field (mean and standard error). The Mito Only and Mito+Rafts preparations contained the same number of donor mitochondria. The 1/10 condition contained 10 % of the mitochondria and 10 % of the lipid rafts used in the Mito+Rafts condition. Significant between group differences were confirmed by single factor ANOVA (F_(3,36)_=31.9; p<0.0001). Post hoc T-tests assuming unequal variances were performed. Significant differences were seen between Mito+Rafts and Mito only, p<0.005; Mito+Rafts and 1/10(Mito+Rafts), p=0.0001; Mito+Rafts and Rafts only, p<0.0001; Mito Only and 1/10(Mito+Rafts), p<0.0005 or Rafts only, p<0.0001; 1/10(Mito+Rafts) and Rafts Only, p<0.001.

To assess whether the transplanted mitochondria were functional, live imaging of cells was performed. For this experiment, donor mitochondria were labeled with Cy-5-mito and recipient cells’ mitochondria were labeled with mitochondria-GFP (green). Cy-5-mito is a far-red fluorescent protein, and was used here instead of red fluorescent DsRed2-mito in order to allow us to have an additional channel available in a non-overlapping wavelength to visualize orange TMRE labeling (574nm). Recipient fibroblasts were incubated with TMRE to assess mitochondrial function after transplantation. TMRE accumulates in mitochondria due to their relative negative charge. Mitochondria that are inactive or depolarized have a decreased membrane potential and therefore do not sequester TMRE. Figure 4 shows (A) internalized mitochondria expressing far-red Cy-5-mito, and (B) part of them appear to be functional (orange TMRE positive), and (C) may have merged with the green GFP recipient mitochondria (or mitochondrial matrix) to some degree, as indicated by (D) the presence of all three fluorescent labels overlapping. The cell nucleus appears blue in image D.

**Figure 4.**
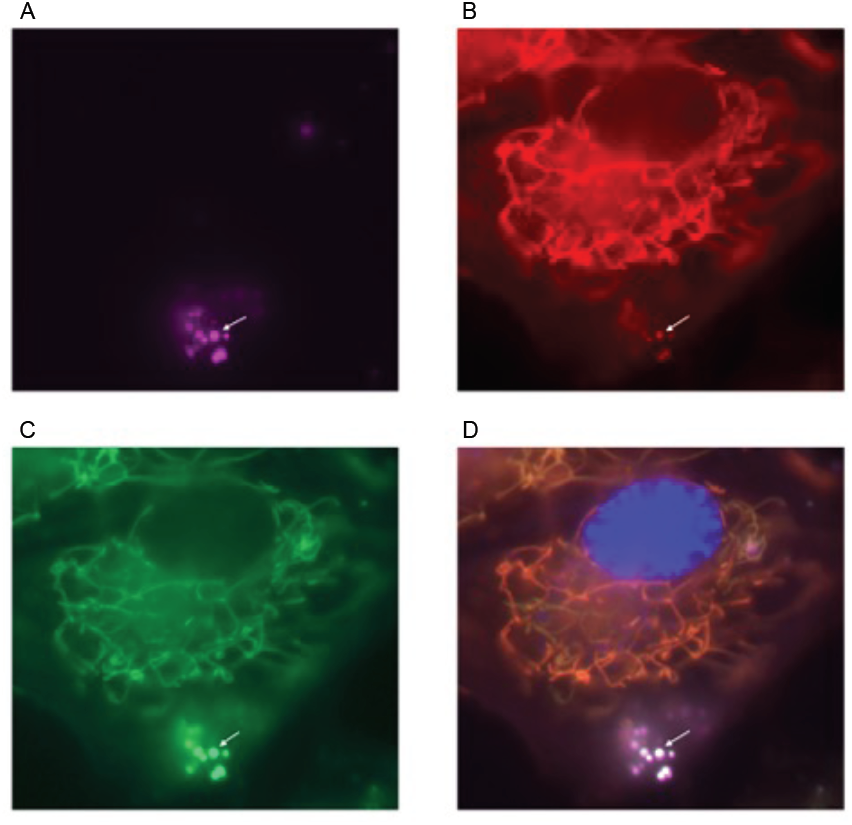
Live imaging of internalized mitochondria indicating that transplanted mitochondria appear functional and can fuse to recipient mitochondria. A) Cy5-mito tagged donor mitochondria. B) TMRE labeled functional mitochondria. C) mitochondria-GFP labeled recipient mitochondria. D) merged image. The arrows indicate an area of potential fusion of functional donor and recipient mitochondria, as evidenced by the overlapping fluorescent labels in each image.

As shown in this figure, donor mitochondria, at the time of measurement, appear largely in the periphery of the recipient cell.

The clinical use of mitochondrial transplantation is currently limited to either fusing donor and recipient cells or injection by needle of mitochondria into recipient cells. Those methods are only suitable for mitochondrial transplantation into individual or small numbers of cells. There are benefits to be gained by exploring and comparing techniques that could be applied to enhance mitochondrial activity in large numbers of cells, as might be used in tissue repair to address mitochondrial deficits that underlie many common disorders, such as disease or aging related dysfunctions of brain and muscle. Several groups have begun experiments with such techniques, in culture, as here, and by injection of mitochondrial preparations into tissue (15,18). The results presented here suggest that for those purposes, exposure of cells to substantially pure mitochondrial preparations packaged in lipids that match cell membranes may produce greater uptake of mitochondria by recipient cells than the use of unpackaged mitochondria. The results do not address the time course of uptake and incorporation of donor mitochondria into the recipient cell and its mitochondrial matrix. Though the mitochondrial matrix is very dynamic, and further movement of elements is expected over time, these studies did not test whether elements from donor mitochondria eventually spread to wide portions of the recipient cell. The results do not address the use of other adjuvants or supplements. They only address syngeneic transplantation, but allogeneic transplantation is also possible. The ideal composition of lipids in the package will require future studies of more alternatives. Nonetheless, the substantial enhancement of uptake seen with lipid packaging suggests that further studies are well worthwhile.

## Author Contributions

BMC conceived and initiated these experiments. BMC, DLM and AMC designed the studies. DLM, LWS and AMC performed the experiments and analyses. DLM and BMC wrote the first draft of the manuscript and BMC, SMB, DLM and DBS made editorial suggestions. Author AMC is deceased. All authors contributed to the manuscript and all living authors have approved the final manuscript.

## Acknowledgements

The studies were performed with the support of private donor funds available to the Program for Neuropsychiatric Research (PI: BMC). No federal grants were obtained nor such grant monies used in the performance of this research. Rachael Neve, Director of the Viral Core Facility at the McGovern Institute for Brain Research at MIT prepared the HSV-DsRed2-mito and HSV-Cy5-mito viruses.

### Competing interests

BMC and AMC are the holders of a patent application for mitochondrial transplantation: “Methods and Compositions for Mitochondrial Replacement," US Appl #12/598,287.

